# Molecular correlates and therapeutic targets in T cell-inflamed versus non-T cell-inflamed tumors across cancer types

**DOI:** 10.1101/773127

**Authors:** Riyue Bao, Jason J. Luke

## Abstract

The T cell-inflamed tumor microenvironment, characterized by CD8 T cells and type I/II interferon transcripts, is an important cancer immunotherapy biomarker. Tumor mutational profile may also dictate response with some oncogenes (i.e. WNT/β-catenin) known to mediate immuno-suppression. Building on these observations we performed a multi-omic analysis of human cancer correlating the T cell-inflamed gene expression signature with the somatic mutanome and transcriptome for different immune phenotypes, by tumor type and across cancers. Strong correlations were noted between mutations in oncogenes and non-T cell-inflamed tumors with examples including *IDH1* and *GNAQ* as well as less well-known genes including *KDM6A, CD11c* and genes with unknown functions. Conversely, we observe many genes associating with the T cell-inflamed phenotype including *VHL* and *PBRM1*, among others. Analyzing gene expression patterns, we identify oncogenic mediators of immune exclusion broadly active across cancer types including HIF1A and MYC. Novel examples from specific tumors include sonic hedgehog signaling in ovarian cancer or hormone signaling and novel transcription factors across multiple tumors. Using network analysis, somatic and transcriptomic events were integrated, demonstrating that most non-T cell-inflamed tumors are influenced by multiple pathways. Validating these analyses, we observe significant inverse relationships between protein levels and the T cell-inflamed gene signature with examples including NRF2 in lung, ERBB2 in urothelial and choriogonadotropin in cervical cancer. Finally, we integrate available databases for drugs that might overcome or augment the identified mechanisms. These results nominate molecular targets and drugs potentially available for immediate translation into clinical trials for patients with cancer.

## Introduction

Despite human cancer being commonly rejected at very early stages, a small number of tumors persist and eventually progress to metastatic disease(Dunn et al. 2002). During the time from tumor formation to systemic spread, genomic and antigenic immuno-editing defines a tumor-immune microenvironment that facilitates the metastatic process(Dunn et al. 2002). Immune profiling of tumor lysates from patients receiving cancer vaccines suggested a paradigm of two broad phenotypes characterized by the presence or absence of CD8+ effector tumor infiltrating lymphocytes (TIL) and other mediators of an adaptive immune response(Harlin et al. 2009). These tumors have been described as T cell-inflamed and demonstrate a transcriptional profile driven by type I interferons as well as expression of many interferon (IFN)-γ linked immunosuppressive mechanisms(Spranger et al. 2013). This is in contrast with non-T cell– inflamed tumors, which have low TIL count and a minimal inflammatory signature. The interaction of the host immunity with tumor cells has been identified as a hallmark of cancer based on the rapid development of immune-checkpoint blocking immunotherapy (Hanahan and Weinberg 2011), predominately centered on programmed-death receptor 1 (PD1)(Ott et al. 2019). While multiple biomarkers have been advanced to better describe the activity of checkpoint immunotherapy, notably tumor mutational burden, a biological predict to an anti-cancer immune response is the elaboration of IFN-γ and the development of T cell-inflamed tumor microenvironment. Given this, the T cell-inflamed phenotype may be leveraged to study factors promoting or limiting a productive anti-tumor immune response.

An aspect impacting efficacy of cancer immunotherapy may be tumor- or tumor microenvironment-intrinsic molecular factors. While immuno-editing may shape the overall antigenic landscape of a tumor, pressure from the immune response will additionally select for tumor clones with specific mutations and/or dysregulation of gene expression pathways that mediate immune evasion. The first tumor-intrinsic oncogene linked to immune-exclusion was β-catenin in the context of metastatic melanoma(Spranger et al. 2015). Activation of the β-catenin pathway, by multiple mechanisms, eliminated required chemokine gradients for the recruitment of BATF3 lineage dendritic cells, with a functional impact of resistance to both checkpoint blocking and adoptive cellular immunotherapy(Spranger et al. 2015; Spranger et al. 2017). Subsequently, this impact of β-catenin has been observed as a mechanism of immuno-suppression across many cancer types(Luke et al. 2019a). A growing list of molecular alterations beyond β-catenin are also being recognized to facilitate immune evasion including PTEN loss or MYC pathway activation (Trujillo et al. 2018). Previous studies have expanded the number of somatic events associated with immune exclusion via computational approaches (Li et al. 2016) (Thorsson et al. 2018) and identified transcriptional signatures that nominate regulators of checkpoint blockage response in patients (Jiang et al. 2018), however have yet to integrate somatic mutations and tumor oncogenic transcriptional pathway activation into the overall model. In addition, it remains unclear whether the resultant immune-suppression of these pathways is due to individual molecular events or whether the phenotype is driven through more complex interactions by multiple signaling pathways.

Using a model wherein the T cell-inflamed tumor microenvironment is considered as a biological approximation of immunotherapy treatment response, we were interested to investigate molecular patterns that associate with the presence or absence of this phenotype with the idea that they may lead to therapeutic opportunities. Here we detail mutations, gene expression patterns and network analyses to nominate molecular aberrations associated with the presence or absence of the T cell-inflamed tumor microenvironment. We further detail available therapeutics that could be explored to reverse or augment this phenotype. These data illuminate the landscape and molecular correlates of the T cell-inflamed tumor microenvironment across cancer types and provide potential immediate next steps for clinical investigation.

## Results

### Candidate cancer oncogenes or tumor suppressors are rarely enriched in most cancer types between T cell-inflamed and non-T cell-inflamed tumors

Using a T cell-inflamed gene expression signature we previously described(Spranger et al. 2016), we categorized tumor samples into non-T cell-inflamed, T cell-inflamed, and intermediate groups across 31 solid tumors from The Cancer Genome Atlas (TCGA). The list of cancer types investigated in this study is provided in **Supplementary Table S1**. To place our approach into clinical context, we correlated our T cell-inflamed gene expression signature with a previously validated interferon-γ associated signature(Ayers et al. 2017) and observed a Pearson’s coefficient of 0.9 (**Supplementary Figure S1**). To investigate the correlation between mutated genes and T cell-inflammation, we investigated 57 candidate cancer oncogenes or tumor suppressors documented in the mutational cancer drivers database IntOGen (05/2016 version) (Tamborero et al. 2018).

For each gene, we compared the frequency of samples harboring non-synonymous somatic mutations (NSSMs) between non-T cell-inflamed and T cell-inflamed groups with relative enrichment of mutated samples shown in **Figure 1A**. Mutations preferentially found in T cell-inflamed tumors included *TP53* (breast cancer), *PIK3CA* (stomach), *BRAF* (colon), *VHL* (Kidney Renal Clear Cell Carcinoma), *CDH1* (stomach), *BRAF* (thyroid), *PTEN* (breast), *KIT* (Uterine Corpus Endometrial Carcinoma), and *ERBB4* (colon) (**Figure 1A**, left panel). In contrast, *FGFR3* (bladder), *TP53* (head and neck, stomach), *NRAS* and *HRAS* (thyroid), and APC (colon) are more frequently mutated in non-T cell-inflamed tumors (**Figure 1A**, right panel). Of the 57 genes investigated, 16 showed relative enrichment of mutated samples in T cell-inflamed or non-T cell-inflamed tumors (**Figure 1B**; P<0.01, unadjusted). No genes passed the significance level of 0.20 after FDR correction for multiple testing effect.

**Figure 1.**
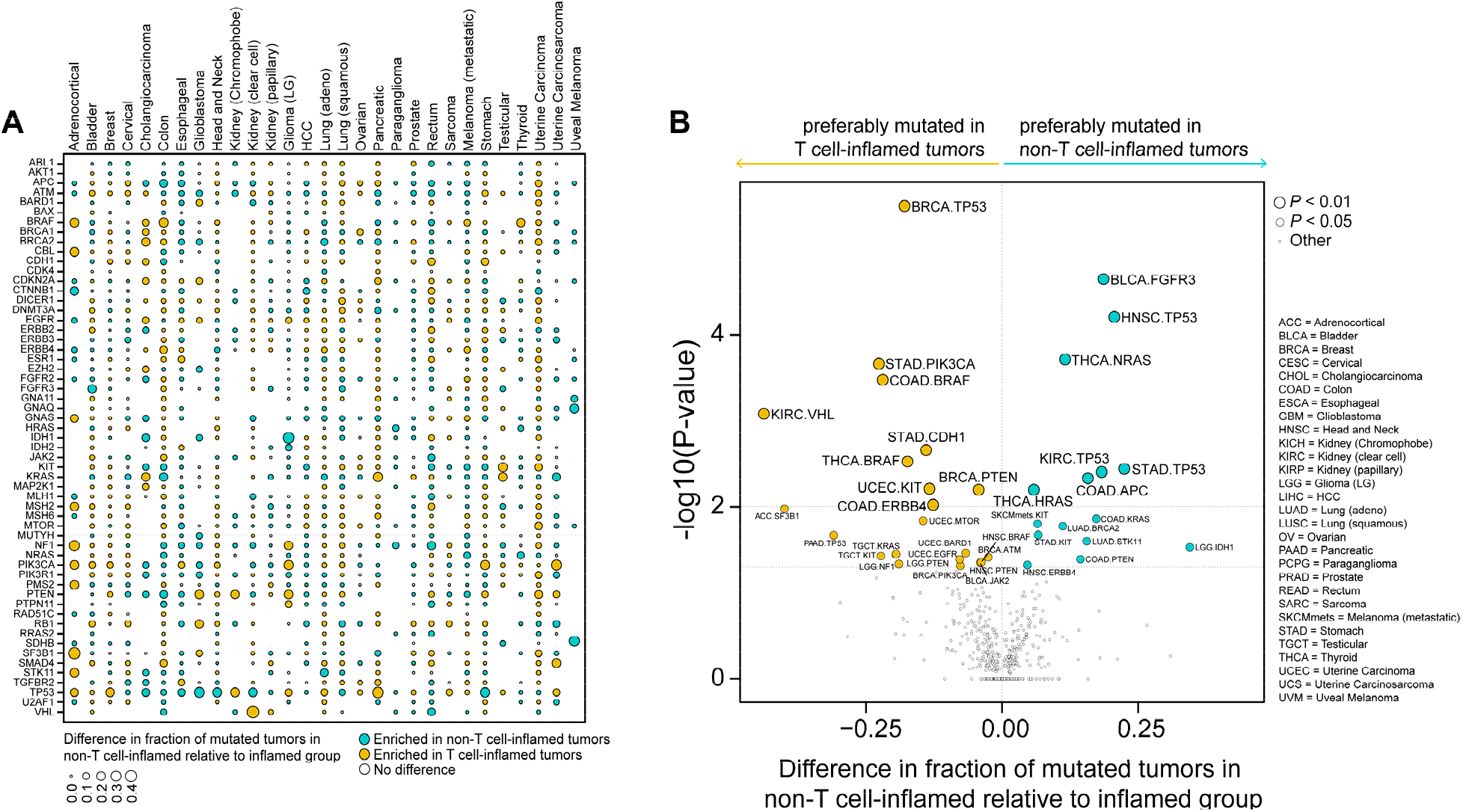
Cancer-specific relative enrichment of mutations in T cell-inflamed and noninflamed tumors from known cancer driver genes. **(A)** 57 gene shown with mutation enrichment in one or more tumor types. Yellow and blue each represents genes preferably mutated in T cell-inflamed and non-inflamed tumor group, respectively. Empty slots indicate this gene is not mutated in this tumor type. **(B)** Cancer-specific mutated genes that are preferably mutated in T cell-inflamed tumors (yellow) or non-inflamed tumors (blue), with the differences in prevalence shown on the x-axis, and p-values shown on the y-axis. Each circle represents one gene in each tumor type. Two-sided Fisher’s exact test was used in **B**.

### Somatic mutations in oncogenic signaling pathways are enriched in non-T cell-inflamed tumors of individual cancer types

To probe molecular mediators of immune-exclusion, we performed an unbiased genome-wide discovery of somatic mutations associated with T cell-inflamed or non-T cell-inflamed tumors within individual cancer types. We collapsed the NSSMs into gene level and counted the number of samples mutated in each of the 18,008 protein-coding genes (see **Methods**). The mutated sample frequency between non-T cell-inflamed and T cell-inflamed groups was built into a contingency table for each gene within each cancer type and compared between groups by twosided Fisher’s exact test. Each tumor type demonstrated a distinct mutation enrichment signature associated with T cell-inflamed or non-T cell-inflamed tumor microenvironment (**Figure 2A**).

Six representative tumor types (papillary thyroid, urothelial, head and neck squamous, clear cell renal cell carcinoma (ccRCC), lung adenocarcinoma and metastatic melanoma) are shown in **Figure 2B**. In urothelial cancer, the gene differentially mutated most frequently in non-T cell-inflamed tumors is *FGFR3*, which was reported in our previous work(Sweis et al. 2016). In head and neck squamous carcinoma, *TP53* mutations are disproportionally mutated between the two groups, with 20.6% more samples mutated in non-T cell-inflamed compared to T cell-inflamed tumors. In lung adenocarcinoma, the gene most significantly mutated was *KEAP1*, recently associated with resistance to PD1 blockade in lung cancer(Rizvi). Alternatively, we identified mutations that are differentially present in T cell-inflamed tumors, such as *PBRM1* in ccRCC, consistent previous reports(Miao et al. 2018; Braun et al. 2019). In metastatic melanoma, *DCTN1* was found to be associated with the presence of T cell-inflammation while *ITGAX* (alias *CD11c)*, is more frequently mutated in non-T cell-inflamed tumors. In papillary thyroid cancer, we detected *BRAF* mutations in at least 17.4% more T cell-inflamed tumors compared to non-T cell-inflamed, and vice versa, *NRAS* and *HRAS* mutations enriched in non-T cell-inflamed phenotype. Data for 23 other tumor types are provided in **Supplementary Figure S2**. Note that the mutations described above passed P<0.01 by two-sided Fisher’s exact test but didn’t reach significance level of 0.20 after FDR correction for multiple testing effect possibly due to sample size. The full list of genes associated with the T cell-inflamed or inflamed phenotype in individual cancer types is provided in **Supplementary Table S2**.

**Figure 2.**
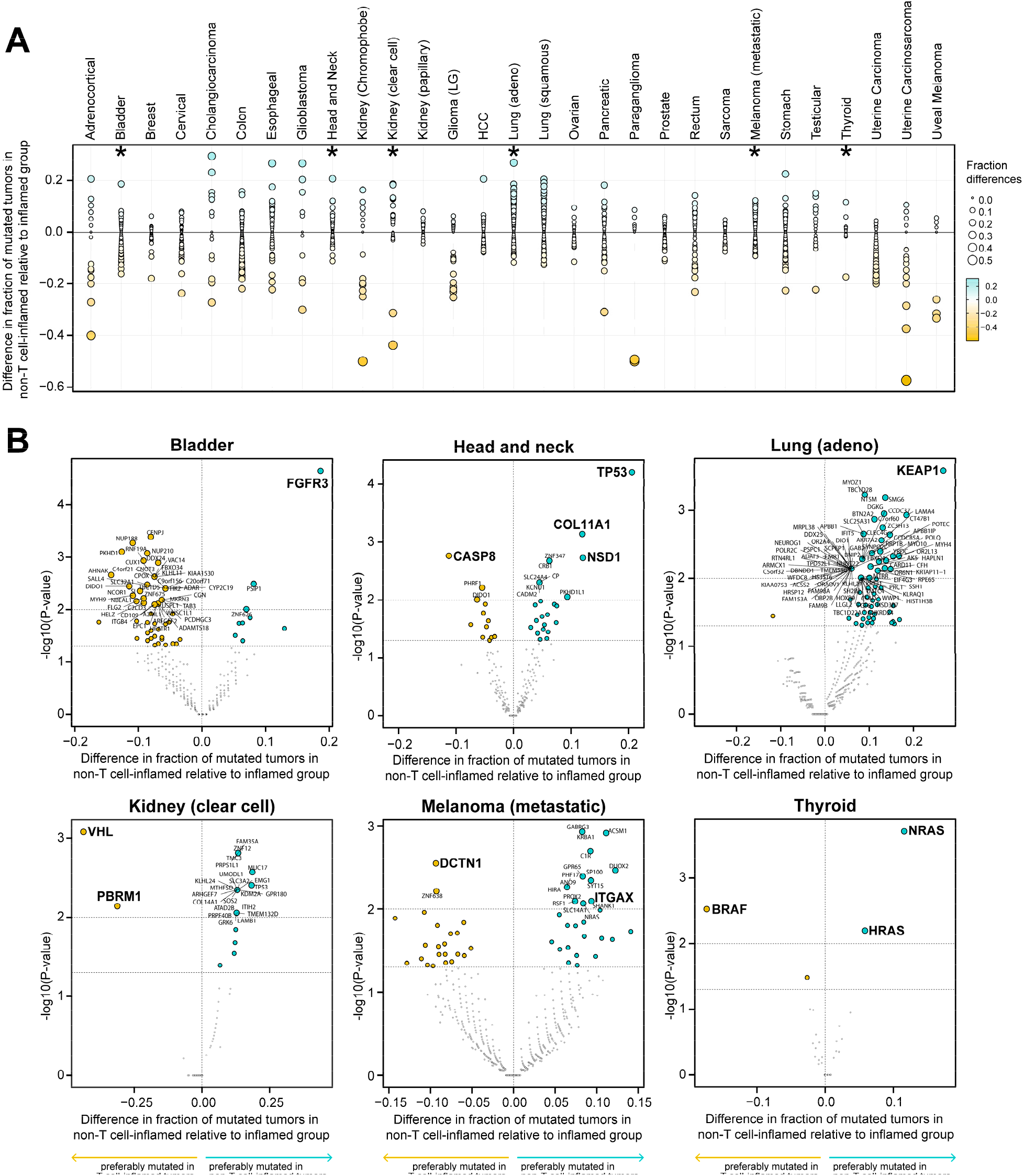
Cancer-specific relative enrichment of mutations in T cell-inflamed and noninflamed tumors from all protein-coding genes on the genome. **(A)** Each tumor type shows a set of gene mutations enriched in T cell-inflamed or non-inflamed tumor group. Stars label the six representative tumor types in **B**, with the rest provided in **Supplementary Figure S2. (B)** Cancer-specific mutated genes that are preferably mutated in T cell-inflamed tumors (yellow) or non-inflamed tumors (blue), with the differences in prevalence shown on the x-axis, and p-values shown on the y-axis. Individual genes of interest discussed in the main text were labeled. Each circle represents one gene in each tumor type. Two-sided Fisher’s exact test was used in **A** and **B**.

### Somatic mutations in oncogenic signaling pathways are shared in non-T cell-inflamed tumors across cancer types

To investigate shared mutations associated with immuno-suppression across multiple tumor types, we performed a pooled analysis of the 29 solid tumors with somatic mutation data available (primary melanoma and mesothelioma didn’t have data available) and identified NSSMs significantly enriched in T cell-inflamed or non-T cell-inflamed tumors. A total of 26 mutated genes were found to be significantly enriched in non-T cell-inflamed tumors at FDR-corrected *P*<0.20 (**Figure 3A**). After categorization by tumor type, the mutations associated with the non-T cell-inflamed phenotype showed a tumor-specific pattern where individual mutations were enriched in some tumor types but not others (**Supplementary Figure S3A**). Mutation in *IDH1* was found to have the strongest association with the non-T cell-inflamed phenotype across all cancers. This mutation was dominantly present in gliomas but also found in other tumor types currently considered non-immunotherapy responsive such as cholangiocarcinoma, breast cancer and sarcoma. Mutation of *GNAQ* is strongly associated with the non-inflamed phenotype across tumors and predominately in uveal melanoma, a subset of melanoma known to be resistant to immune-checkpoint blockade(Luke et al. 2013). Several of the identified genes are highly associated with the non-T cell-inflamed phenotype across cancers and regulate shared or connected signaling pathways. For example, *CTNNB1, APC*, and *AXIN1* from the β-catenin pathway cross-talk with PTEN signaling(Lague et al. 2008) and all have been associated with immune exclusion in our prior work(Luke et al. 2019b). Mutation in *KDM6A* has not previously been described as mediating the non-T cell-inflamed phenotype however KDM6A is a known epigenetic regulator of interleukin-6 and IFN-β(Li et al. 2017) and linked to autoimmunity (Ming et al. 2005).

We used a more stringent significance level cutoff (FDR-corrected *P*<0.05) for the genes enriched in the T cell-inflamed phenotype (**Figure 3B**). In contrast with non-T cell-inflamed enriched mutations, mutations associated with the T cell-inflamed phenotype demonstrated a more homogenous distribution across tumor types (**Supplementary Figure S3B**). Among the most statistically significant were *VHL* and *PBRM1*, which was found by a second method in pancancer analysis (FDR-corrected P= 8.08e-14). In overview, **Figure 3C** demonstrates that a larger number of genes were found to be associated with the presence of T cell-inflammation from the pan-cancer analysis, with a distinct set of genes more frequently mutated in non-T cell-inflamed tumor group. The full list of genes associated with the T cell-inflamed or non-T cell-inflamed phenotype in pan-cancer analysis is provided in **Supplementary Table S3**.

**Figure 3.**
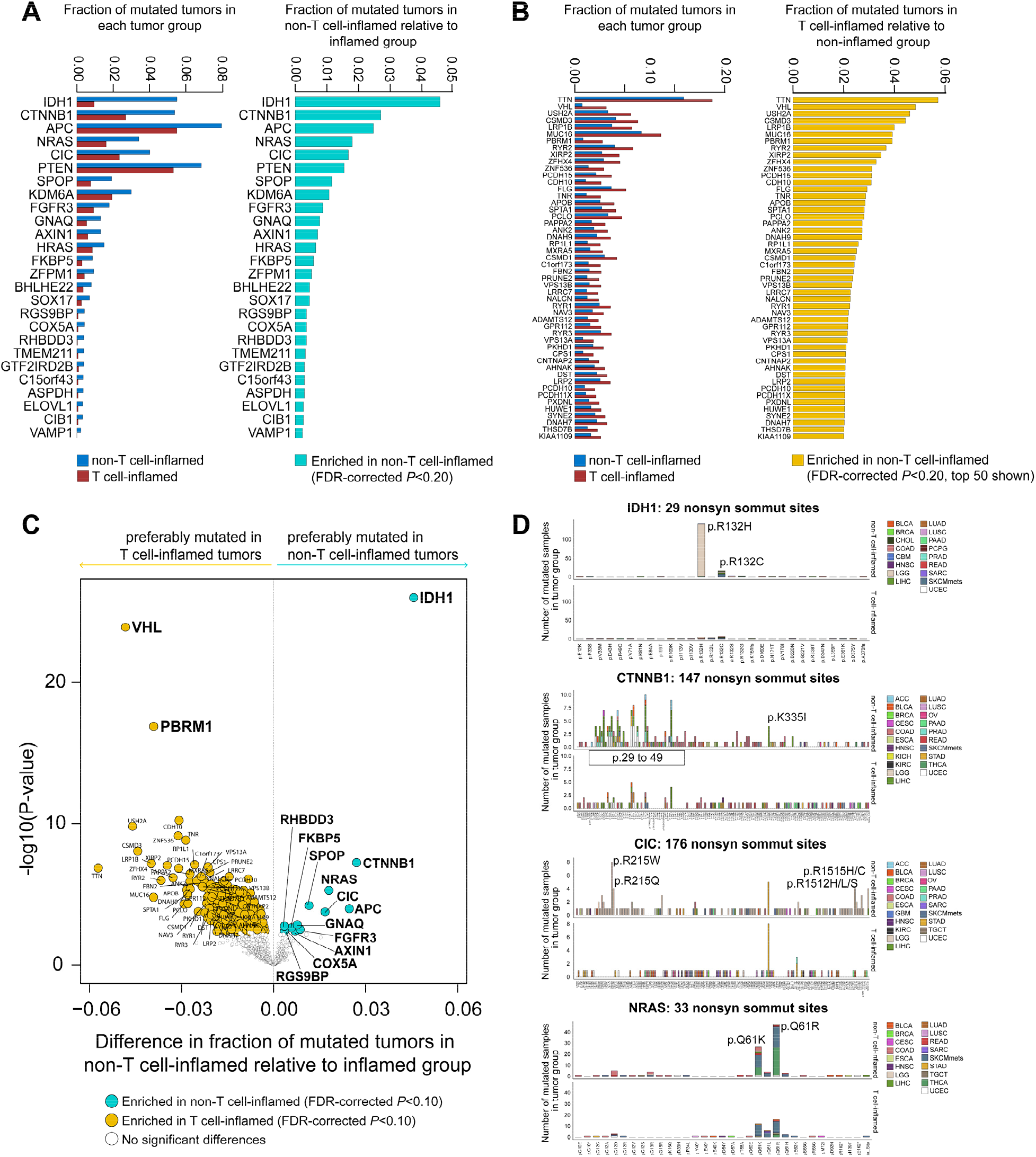
Pan-cancer relative enrichment of mutations in T cell-inflamed and non-inflamed tumors from all protein-coding genes on the genome. **(A)** 26 genes carrying NSSMs more frequent in non-T cell-inflamed relative to inflamed tumors at FDR-corrected *P*<0.20. **(B)** Top 50 genes carrying NSSMs more frequent in T cell-inflamed relative to non-inflamed tumors at FDR-corrected *P*<0.05. **(C)** Distribution of relative enrichment of mutated tumors in T cell-inflamed or non-inflamed tumor group. Gene symbols labeled represent those more frequently mutated in T cell-inflamed (yellow) or non-T cell-inflamed (cyan) at FDR-corrected *P*<0.10. A different p-value cutoff was selected here to allow readability of the gene symbols on figure. **(D)** Representative examples showing the protein-domain-specific enrichment of non-T cell-inflamed mutations in *IDH1, CTNNB1, CIC* and *NRAS* pan-cancer wide. Two-sided Fisher’s exact test was used in AD, with FDR multiple testing correction.

### Activation of transcriptional programs correlates with the non-T cell-inflamed tumor microenvironment across cancer types

Our previous work suggested that activation of transcriptional programs may be another mechanism associated with the non-T cell-inflamed phenotype and only partially overlaps with somatic mutations(Luke et al. 2019a). Therefore, we performed unbiased genome-wide pathway discovery following the same protocol as previously described (Luke et al. 2019a). Differentially expressed genes (DEGs) were identified between non-T cell-inflamed and T cell-inflamed tumors within each cancer type, and upstream regulators were predicted based the DEG-encoding target molecules using causal network analysis (Kramer et al. 2014). Pathways were ranked by the number of tumor types sharing activation of the same pathway (**Figure 4A**; all pathways shown in **Supplementary Figure S4**, and their activation status in individual cancer type provided in **Supplementary Table S4**).

Among the top 30 pathways, we observed a strong impact of *CTNNB1* as well as several other pathways described in the literature as influencing anti-tumor immunity such as *MYC* (Kortlever et al. 2017), *PPARG* (Korpal et al. 2017) and *SHH* (sonic hedgehog) (Shi et al. 2018). In addition, we identified transcriptional programs correlated with the non-T cell-inflamed phenotype such as *KLF4, HIF1A, KMT2D (alias MLL2), NFE2L2 (alias NRF2)*, androgen receptor (AR), and estrogen receptor (ER) signaling, which have some literature support for modulating the immune response(Rizvi; Kovats 2015; Ou et al. 2019).

Cognizant of crosstalk between canonical signaling pathways, we sought to investigate the connections between activated transcriptional programs as related to the non-T cell-inflamed tumor phenotype. To pursue this, upstream regulator encoding genes were annotated with KEGG pathways, and built into a network with nodes representing each KEGG pathway and edges representing the shared molecules between pathways. KEGG pathways were observed to be highly connected including PI3K-Akt signaling, MAPK signaling, and FoxO (Forkhead Box O) signaling (**Figure 4B**).

To validate the activation of transcriptional programs at protein level, we correlated the normalized RPPA protein abundance of the predicted upstream regulator and/or aggregation of the downstream target molecules with the T cell-inflamed gene expression signature for each tumor type. Out of 266 activated pathways, 60 (22.6%) have corresponding proteins measured on the RPPA platform and were included in the correlation analysis (**Figure 4C**). Among these, 54 (90.0%) showed anti-correlation with the T cell-inflamed gene expression signature (Pearson’s correlation coefficient r<0). Twenty-seven (45.0%) pathways reached significance level of 0.10 after FDR-correction for multiple testing. Representative pathways include NRG1 in testis cancer, NFE2L2 *(alias* NRF2) in lung squamous cancer, ERBB2 (*alias* HER2) in bladder cancer, and Cg in ovarian cancer (**Figure 4D**). Complete listing of RPPA correlations are found in **Supplementary Data File S1**.

**Figure 4.**
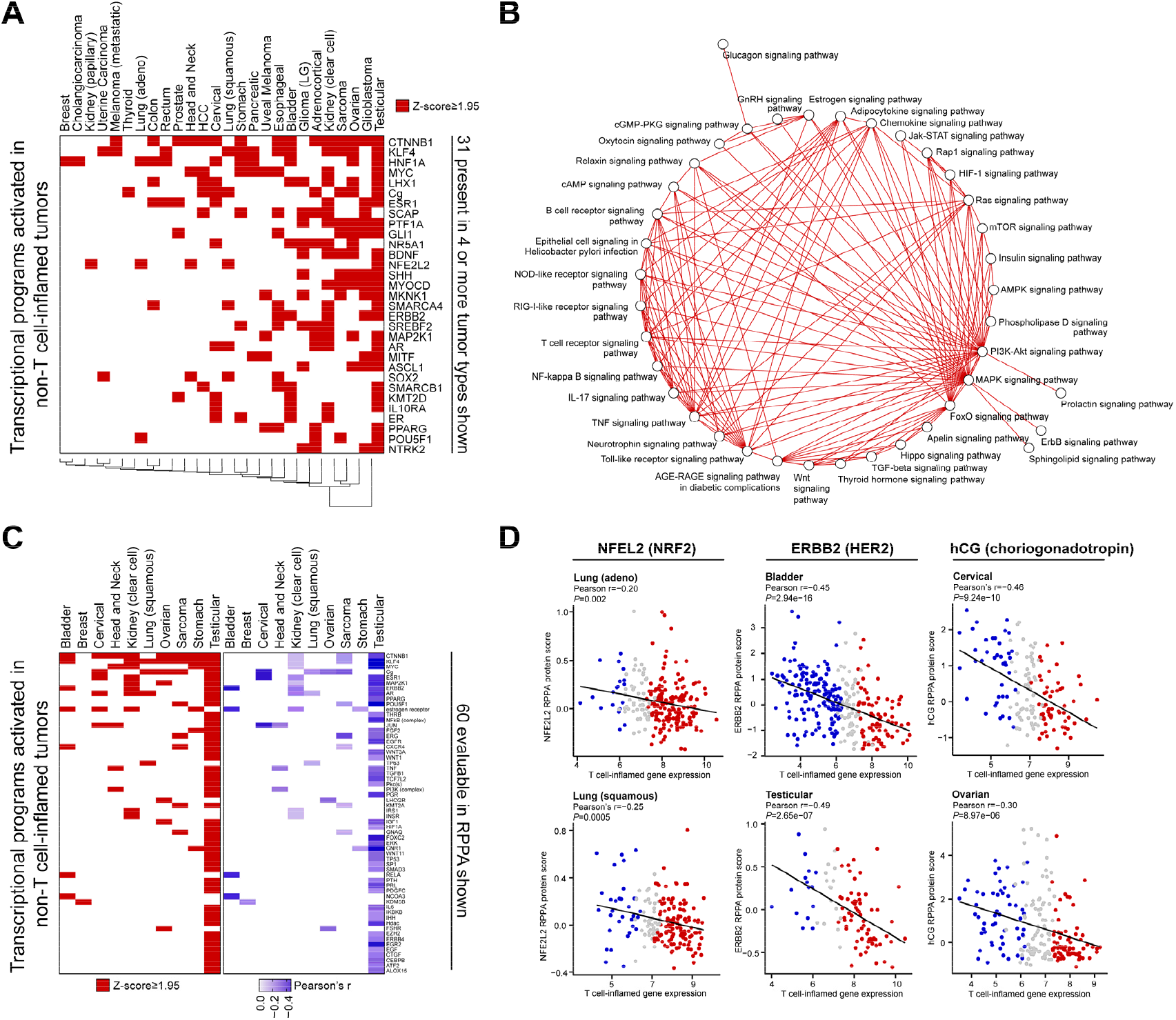
Pan-cancer pathway activation in non-T cell-inflamed relative to inflamed tumors from RNAseq gene expression. **(A)** Each tumor type shows transcriptional programs activated in non-T cell-inflamed relative to the inflamed tumor group. 31 transcriptional programs present in 4 or more tumor types are shown. Red color labels those at activation z-score ≥ 1.95 predicted by IPA causal network analysis (see Methods). **(B)** Pathway analysis shows connections between multiple transcriptional programs activated in non-T cell-inflamed tumors from one or more type types. **(C)** RPPA validation of activated transcriptional programs at protein level. 60 transcriptional programs evaluable in RPPA data are shown. Left panel: activation score heatmap of the 60 transcriptional programs in non-T cell-inflamed relative to inflamed tumors in each tumor type, with red color indicating z-score ≥ 1.95; right panel: Pearson’s correlation coefficient heatmap of each transcriptional program vs. T cell-inflamed gene expression signature, with blue color indicating negative correlation (Pearson’s r < 0). **(D)** Representative correlation between activated upstream regulator NFEL2 *(alias* NRF2), ERBB2 (*alias* HER2), hCG and T cell-inflamed gene expression signature. Full results are provided in **Supplementary Data File S1**. IPA’s causal network analysis is used in **A**, Pearson’s correlation is used in **C** and **D**.

### Mutations or pathways associated with either the T cell-inflamed or non-T cell-inflamed phenotype co-occur within individual tumors

To investigate the interaction between somatic mutations and/or activated transcriptional programs, we summarized the number of mutations or activated pathways on the per-patient level. Out of 3,137 patients from the non-T cell-inflamed tumor group, 1,062 (33.8%) harbor NSSMs (**Figure 5A**) in at least one of the 26 non-T cell-inflamed genes from our previous analysis above and 2,259 (72.0%) show activation in at least one of the 31 transcriptional programs from those reported above (**Figure 5B**). Of the 1,062 patients, 363 (34.2%) harbor NSSMs in two or more genes (**Figure 5C**) and 1,870 out of 2,259 patients (82.8%) demonstrate activation in two or more transcriptional programs (**Figure 5D**). Taking together, 2,423 patients (77.2%) across cancer types demonstrate evidence of at least one resistance mechanism by mutation or transcriptional program activation (**Supplementary Table S5**).

To facilitate clinical translation of these results, we were interested to identify existing drugs that might be repurposed as immunomodulatory agents. To pursue this, we queried an existing United States Food and Drug Administration (FDA) ensemble database. The majority of the relevant molecular mechanisms identified in our study were found to have one or more potentially relevant drugs available (**Supplementary Figure S5; Supplementary Table S6**). Inhibitors were identified for mutated genes and signaling pathways activated in non-T cell-inflamed tumors, such as FGFR3, HRAS, IDH1, NRAS, PTEN, AR, MAP2K1 and NTRK2 which may nominate therapeutic approaches to overcome the non-T cell-inflamed phenotype. Alternatively, well known agents targeting IFNG, IL2 or TLR9 might be considered as adjuncts in tumor types already demonstrating the T cell-inflamed phenotype.

**Figure 5.**
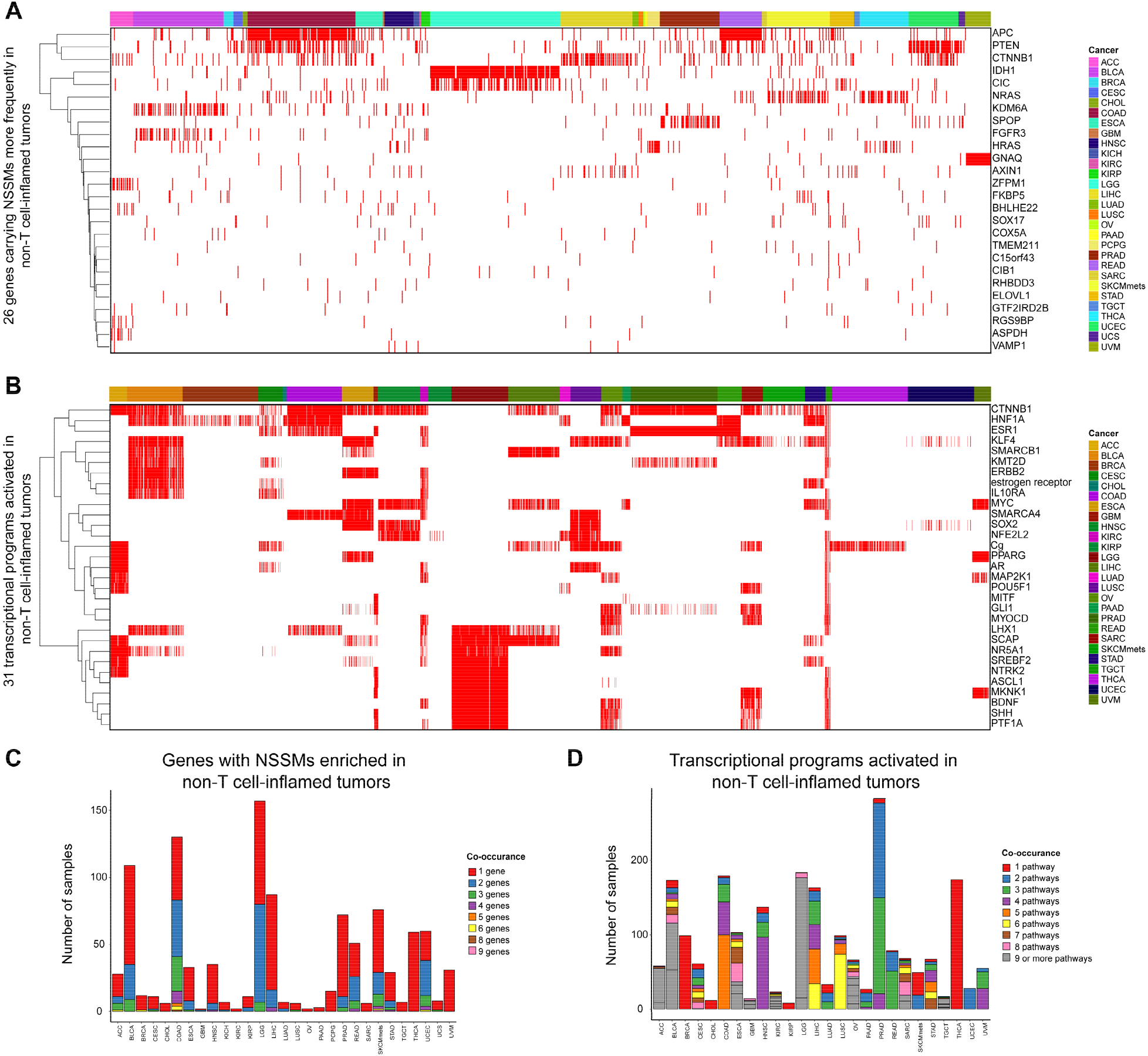
Multiple mutation or pathway-mediated resistance mechanisms co-occur in non-T cell-inflamed tumors. **(A)** Co-occurrence of NSSMs in 26 genes from **Figure 3A** at per-patient level. Annotation bar above the heatmap indicates tumor type. **(B)** Co-activation of 31 transcriptional programs from **Figure 4A** at per-patient level. Annotation bar above the heatmap indicates tumor type. **(C)** Distribution of tumor samples with one or more genes carrying NSSMs in the same patient from non-T cell-inflamed tumor group. Eight categories are shown; category of co-occurrence in 7 genes is not shown because no sample fell into this category. **(D)** Distribution of tumor samples with one or more transcriptional programs activated in the same patient from non-T cell-inflamed tumor group. Nine categories are shown.

## Discussion

The T cell-inflamed tumor microenvironment, characterized by TIL and evidence for IFN-γ linked adaptive immunity, has been strongly associated with clinical response to checkpoint immunotherapy(Ayers et al. 2017; Cristescu et al. 2018). Based on our prior observations surrounding the impact of dysregulated oncogenic signaling from the WNT/β-catenin pathway, we performed a per tumor type and pan-cancer analysis to assess the landscape of somatic and transcriptomic events associated with the T cell-inflamed or non-T cell-inflamed phenotype. We stratified solid tumors of TCGA by inflamed, intermediate and non-inflamed status and interrogated genomic aberrations as well as gene expression patterns associated with these phenotypes. We identified groups of somatic mutations strongly linked to T cell-inflamed or non-T cell-inflamed tumors including some in well-known genes and as well as others that are less well understood. Similarly, we have identified pathways associated with these phenotypes with some established as immuno-suppressive and others previously unknown.

A growing list of dysregulated tumor-intrinsic oncogenes have been described which mediate immune exclusion by multiple mechanisms. Taken together these somatic events broadly lead to a state of decreased innate immunity, antigen presentation and T cell trafficking to tumors. For example mutation of *STK11* has been strongly associated with lack of response to checkpoint inhibitors(Skoulidis et al. 2018) and mechanistic studies have linked this aberration to suppression of STING agonism(Kitajima et al. 2019). Similarly MYC activation is known to be reversibly associated with lack of response to immunotherapy(Topper et al. 2017) and hypoxic environments(Gnanaprakasam et al. 2017), commonly associated with liver metastases which are less responsive to immunotherapy(Johnson et al. 2019).

In our cancer type specific and pan-cancer analysis we observed the presence of several mutations that deserve further interrogation on a mechanistic level. For example the histone demethylase encoded by KDM6A, where gene deletion has long been associated with the Kabuki immunodeficiency syndrome(Ng et al. 2010), is also known to be associated with immune relevant processes including the type I interferon(Li et al. 2017) and hypoxia response(Chakraborty et al. 2019). In metastatic melanoma we observed *ITGAX(alias CD11c)*, a marker expressed on myeloid cells with well-known impact on classical and monocytic dendritic cell function(Wculek et al. 2019), is more frequently mutated in non-T cell-inflamed tumors.

As opposed to tumor-intrinsic oncogenic mutations driving immuno-suppression, we have additionally identified recurrent somatic events that may potentiate the anti-tumor immune response. The most obvious of these, *PBRM1*, has previously been shown from patient samples in renal cell carcinoma to impact outcomes to checkpoint blockade (Miao et al. 2018) and has been demonstrated to augment T cell-mediated killing(Pan et al. 2018). We have identified a group of further mutations that should be explored for functional validation and analyzed from patient tumor databases. An intriguing consideration surrounding both immuno-suppressive and immuno-activating mutations is the possibility of somatically engineering such mutations into tumor cells prior to the initiation of immunotherapy through DNA modifying approaches such as oncolytic viruses(Emdad et al. 2018) or *in vivo* CRISPR(Chow and Chen 2018).

In addition to somatic mutations associated with immune phenotypes, our transcriptional analysis of gene programs associated with T cell-inflamed and non-T cell-inflamed tumors has nominated multiple immediate and novel therapeutic approaches. Some of these such as hedgehog signaling in ovarian cancer and hormone receptor (estrogen and androgen) in multiple diseases including urothelial or adrenocortical cancers deserve investigation as combination approaches with immunotherapy. In this setting however it would be important to assess for transcriptional activation to confirm that the hypothesis was investigated in the correct patient population.

While we and others have previously described transcriptional patterns in individual diseases linked to immuno-suppression, our data in this analysis argues for transcriptional programs that are substantially shared across tumor types. We see CTNNB1, KLF4, HIF1A, and MYC as particularly dominate across many cancers as well as evidence to suggest substantial cross talk and perhaps overlap. Because of this we were interested to investigate the interactions of these pathways and noted MYC signaling as perhaps the most important node associating with lack of the T cell-inflamed phenotype. Consideration then of approaches to alleviate MYC related immuno-suppression should be a high priority.

While targeting of some oncogenic pathways remains difficult, our analysis has identified multiple targets for which translation is possible with immediate examples surrounding the PI3K, MAPK and FGFR pathways. Beyond these we have particularly focused on IDH1 as a metabolism mediator of immuno-suppression. Based on our data in concert with functional validation from human and murine models(Kohanbash et al. 2017), we have taken forward a phase II clinical trial of the IDH1 inhibitor ivosidenib with anti-PD1 in patients selected by *IDH1* mutation (NCT04056910). More broadly, our FDA database analysis suggests that there are many drugs with the potential to be re-evaluated, re-purposed as immune modifiers potentially to combine with immunotherapy.

Noting that some of the findings from our analysis already have confirmation from patient tissue and clinical trials, we do acknowledge limitations of this work. Using the T cell-inflamed gene expression phenotype is an imperfect approximation of patient response in clinical trials however interferon associated gene expression does appear to perhaps be the best translational biomarker currently available after failure of TMB in multiple phase III trials. Additionally, our analysis of TCGA does not facilitate specific outcomes to immunotherapy treatment response however the size of the dataset allows a large discovery power to identify novel genes and pathways associated with immune exclusion.

Treatment with cancer immunotherapy has become a backbone of cancer therapy however is ineffective in the majority of patients and novel biomarkers as well as molecular targets are needed to advance the field. In an unbiased analysis, we have identified molecular alterations already validated from patient tissues but more importantly many somatic and transcriptional aberrations that deserve immediate study. Based on these data, we have already advanced to clinical trials and believe that many drugs are available in the public domain that might be considered for immunotherapy re-purposing. As other biomarkers or novel insights (i.e. – epigenetics) become more robust it may be possible to integrate this data into similar analyses to identify further mediators of immunotherapy response and resistance.

## Methods

### TCGA cancer datasets

Level 3 RNA-seq gene expression (release February 4, 2015), level 4 somatic mutations (release January 28, 2016), and clinical tables (release January 28, 2016) were downloaded for 31 solid tumor types from The Cancer Genome Atlas (TCGA) preprocessed by Broad’s team (https://gdac.broadinstitute.org/). Survival data consisting of four major clinical endpoints were obtained from standardized clinicopathologic annotation data tables in TCGA Pan-Cancer Clinical Data Resource (TCGA-CDR) (Liu et al. 2018). Gene expression was quantified by Expectation Maximization (RSEM) algorithm (Li and Dewey 2011), with raw read counts mapped to gene features and transcripts per million (TPM) reported. The count-based gene expression was normalized across all samples using upper quartile method followed by log_2_ transformation. A total of 9,555 tumor samples were initially downloaded. After excluding three tumor types consisting of high tumor cell-intrinsic immune cell transcripts (acute myeloid leukemia, diffuse large B-cell lymphoma, and thymoma), 9,244 tumors were included in the analysis.

### Identification of non-T cell-inflamed and T cell-inflamed tumor groups

The tumor samples were grouped into non-T cell-inflamed, T cell-inflamed, and intermediate categories using a defined T cell-inflamed gene expression signature consisting of 160 genes same as previously described (Spranger et al. 2016). In brief, the normalized gene expression was transformed into a scoring system where each gene is defined as upregulated (+1), downregulated (−1), or unchanged (0) relative to the median of its expression across all samples. By adding scores of all 160 genes from the signature together, we obtained a sample-wise gene score ranging from −160 (if all genes are downregulated) to +160 (if all genes are upregulated). Tumors of score < −80 were groups as non-T cell-inflamed, those of score > +80 as T-cell inflamed, and the rest as intermediate.

### Differential gene expression detection and pathway activation prediction from RNAseq data

Within each individual cancer type, the non-T cell-inflamed group was contrasted against the T cell-inflamed group and differentially expressed genes (DEGs) were identified by Linear Models for Microarray and RNA-Seq Data (limma) voom algorithm with precision weights (v3.38.3) (Law et al. 2014). The empirical quality weights were estimated to account for variation in precision among samples and were shown to increase performance of DEG detection especially for highly heterogeneous human tumor samples. Significant DEGs were filtered by FDR-adjusted *P*<0.05, and fold change ≤ 1.5 or ≥ −1.5. Pathways significantly affected by the DEGs were detected by IPA^®^ (QIAGEN Inc., Germany) with the curated Ingenuity Knowledge Base (accessed November 2015). Upstream transcriptional regulators and their change of direction were predicted based on the cumulative effect of downstream target molecules (encoded by DEGs) implemented in IPA® casual network analysis (Kramer et al. 2014). Those of overlap *P*<0.05 (measuring the enrichment of target molecules in the dataset) and activation z-score>1.95 were selected for further analysis. The predicted upstream regulators were further annotated by Kyoto Encyclopedia of Genes and Genomes (KEGG) database and used to build a pathway-to-pathway network based on shared upstream regulators among annotated pathways in Cytoscape (v3.7.1), with nodes denoted as individual KEGG signaling pathways, and edge size denoted as the number of shared upstream regulators between pathways.

### Differential somatic mutation enrichment detection from whole exome sequencing data

For somatic mutations, those predicted to be affecting protein-changing sequences (referred to as, NSSMs, non-synonymous somatic mutations), including non-synonymous/stoploss/stopgain small nucleotide variants (SNVs), frameshift/non-frameshift small insertions and deletions (indels), and variants that affect the splicing site, were kept for analysis. The NSSMs were summarized into gene level, and then into sample level to generate the number of samples carrying mutations in this gene (referred to as, mutated samples), with the rest of the samples categorized as non-mutated. The frequency of gene-wise mutated or non-mutated samples was compared between non-T cell-inflamed and T cell-inflamed tumor group using twosided Fisher’s exact test. Genes that are more frequently mutated in non-T cell-inflamed or T cell-inflamed group were defined as non-T cell-inflamed and T cell-inflamed mutated genes, respectively. Pathways significantly affected in non-T cell-inflamed or non-T cell-inflamed-enriched mutated genes were identified by IPA® (QIAGEN Inc., Germany).

### RPPA data analysis

Level 3 reverse phase protein array (RPPA) antibody-level protein abundance data (release January 28, 2016; patch July 14, 2016) produced by MD Anderson Cancer Center were downloaded from TCGA preprocessed by Broad’s team (https://gdac.broadinstitute.org/) (accessed April 26, 2018). For each protein, its abundance was estimated using median-centered normalized values corresponding to the antibody from the data files without gene annotation. Within each tumor type, a one-sided Pearson’s correlation (R function *contest*, alternative = “less”) test was performed within each tumor type between the abundance of a protein and T-cell inflamed gene expression from normalized RNA-seq data, followed by FDR-correction for multiple testing.

### Drug-gene interaction and druggable gene category identification

Candidate genes identified by pathway activation or somatic mutation enrichment analysis were queried in The Drug Gene Interaction Database (Cotto et al. 2018) (DGIdb, http://www.dgidb.org) (v3.0, accessed February 25, 2019) for gene-drug interactions (inhibitor, activator, etc.) and druggable gene categories (clinically actionable, drug resistance, etc.) with default settings. The DGIdb integrates resources from 20 databases and consists of existing drugs including those that are FDA-approved, antineoplastic, and/or immunotherapies.

### Statistical analysis

For analysis of contingency tables including comparison of sample frequency between groups activated by pathways, two-sided Fisher’s exact test was used. Gene expression comparison between tumor groups were performed using linear regression models implemented in limma voom with precision weights. For multiple comparisons, p-value was adjusted using Benjamini-Hochberg FDR correction for multiple testing (Benjamini et al. 2001). Pearson’s correlation r was used for measuring statistical dependence between normalized and log2-transformed expression level of different genes. Statistical analysis was performed using R (v3.5.1) and Bioconductor.

### Data access

The original data files were downloaded from TCGA and publicly available on Broad’s GDAC-Firehose website (http://gdac.broadinstitute.org/). Preprocessed data files are provided as supplementary tables. Additional large-size preprocessed data files will be provided upon request from corresponding author.

