## Supplementary Figures for "Molecular correlates and therapeutic targets in T cell-inflamed versus non-T cell-inflamed tumors across cancer types"

**Running Title:** Molecular correlates of inflamed or non-inflamed tumors

**Authors**: Riyue Bao[1], Jason J. Luke[1,*]

[1]University of Pittsburgh Medical Center, Pittsburgh, PA

[*]**Corresponding Author:**
Jason J. Luke, MD, FACP
Associate Professor of Medicine
University of Pittsburgh Medical Center and Hillman Cancer Center
5150 Centre Ave. Room 564
Pittsburgh PA 15232


**Keywords**: T cell-inflamed, immune evasion, genomics, transcriptomics, TCGA

**Support**: JJL acknowledges Department of Defense Career Development Award (W81XWH-17-1-0265), the Arthur J Schreiner Family Melanoma Research Fund, the J. Edward Mahoney Foundation Research Fund, Brush Family Immunotherapy Research Fund and Buffet Fund for Cancer Immunotherapy.

**Conflict of Interest Disclosures**: RB: None; JJL declares Data and Safety Monitoring Board: TTC Oncology, Scientific Advisory Board: 7 Hills, Actym, Alphamab Oncology, Array, BeneVir, Mavu, Tempest, Consultancy: Aduro, Astellas, AstraZeneca, Bayer, Bristol-Myers Squibb, Castle, CheckMate, Compugen, EMD Serono, IDEAYA, Immunocore, Janssen, Jounce, Leap, Merck, Mersana, NewLink, Novartis, RefleXion, Spring Bank, Syndax, Tempest, Vividion, WntRx, Research Support: (all to institution for clinical trials unless noted) AbbVie, Array (Scientific Research Agreement; SRA), Boston Biomedical, Bristol-Myers Squibb, Celldex, CheckMate (SRA), Compugen, Corvus, EMD Serono, Evelo (SRA), Delcath, Five Prime, FLX Bio, Genentech, Immunocore, Incyte, Leap, MedImmune, Macrogenics, Novartis, Pharmacyclics, Palleon (SRA), Merck, Tesaro, Xencor, Travel: Array, AstraZeneca, Bayer, BeneVir, Bristol-Myers Squibb, Castle, CheckMate, EMD Serono, IDEAYA, Immunocore, Janssen, Jounce, Merck, Mersana, NewLink, Novartis, RefleXion, Patents: (both provisional) Serial #15/612,657

### Supplementary Figures

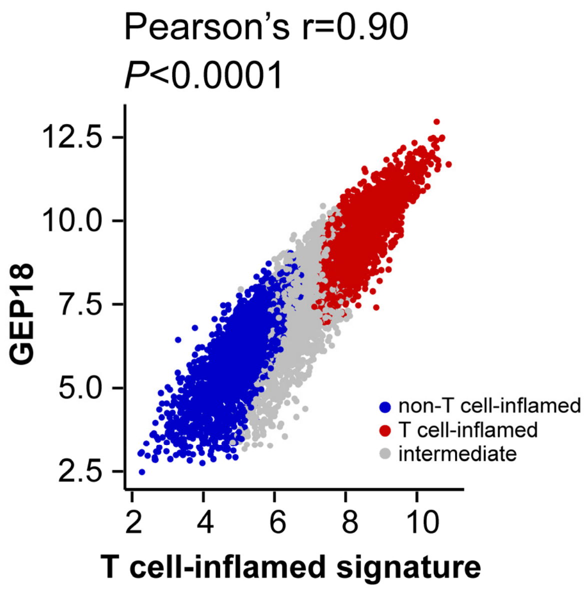


**Supplementary Figure S1. T cell-inflamed gene expression signature is highly correlated with previously published GEP18 signature.**

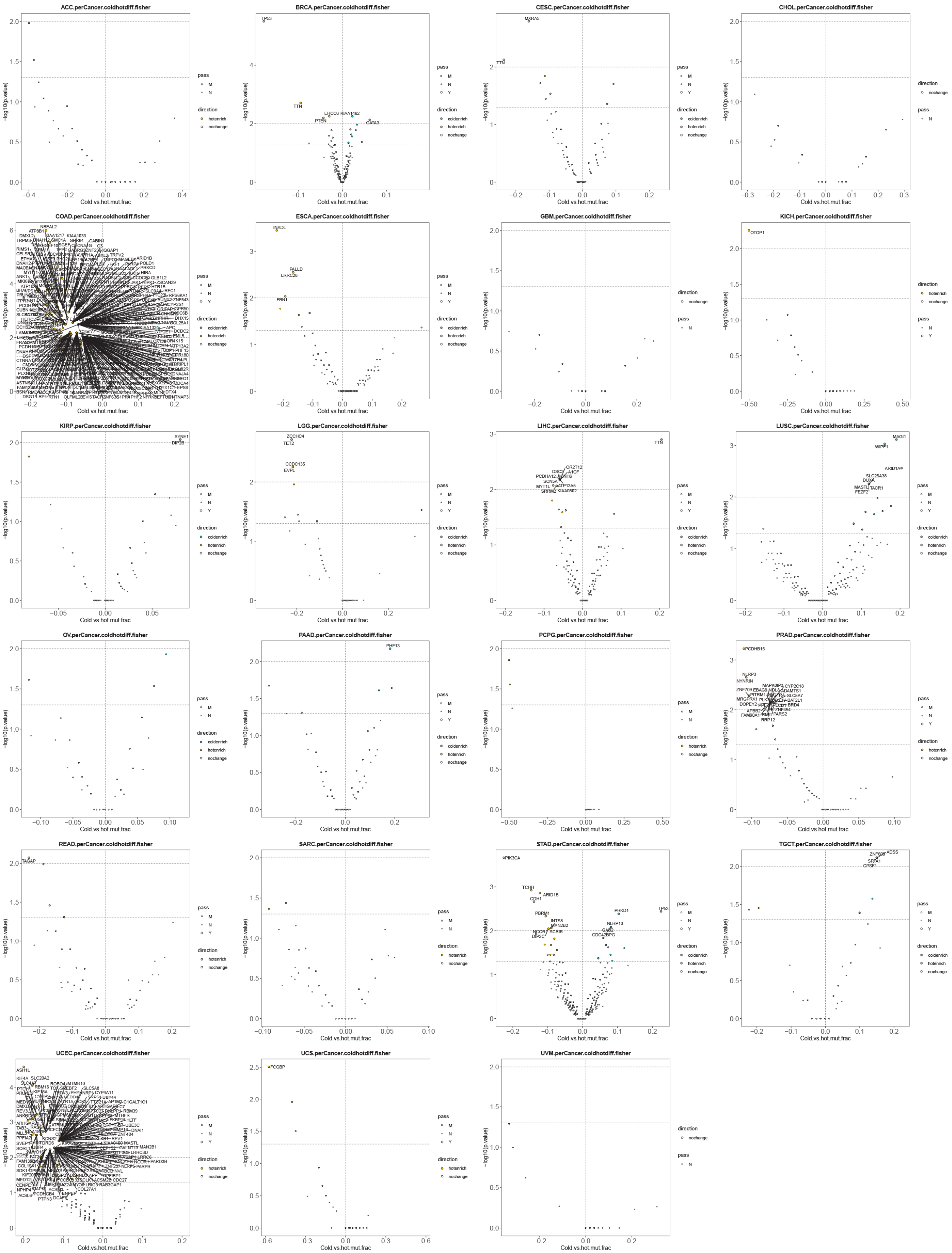
ACC.perCancer.coldhotdiff.fisher
BRCA.perCancer.coldhotdiff.fisher
TP53
TTN
ERCC6
KIAA1462
PTEN
GATA3
CESC.perCancer.coldhotdiff.fisher
MXRA5
TTN
CHOL.perCancer.coldhotdiff.fisher
COAD.perCancer.coldhotdiff.fisher
ESCA.perCancer.coldhotdiff.fisher
INADL
PALLD
LRRC2B
FBN1
GBM.perCancer.coldhotdiff.fisher
KICH.perCancer.coldhotdiff.fisher
OTOP1
KIRP.perCancer.coldhotdiff.fisher
SYNE1
DIP2B
LGG.perCancer.coldhotdiff.fisher
ZCCHC4
TET2
CCDC135
EVPL
LIHC.perCancer.coldhotdiff.fisher
TTN
LUSC.perCancer.coldhotdiff.fisher
MAGI1
WIPF1
ARID1A
SLC25A38
DUXA
MASTL
TACR1
FEZF2
OV.perCancer.coldhotdiff.fisher
PAAD.perCancer.coldhotdiff.fisher
PHF13
PCPG.perCancer.coldhotdiff.fisher
PRAD.perCancer.coldhotdiff.fisher
PCDHB15
READ.perCancer.coldhotdiff.fisher
TAGAP
SARC.perCancer.coldhotdiff.fisher
STAD.perCancer.coldhotdiff.fisher
PIK3CA
TCHH
ARID1B
CDH1
PBRM1
INTS8
MAN2B2
NCOR1
SCRIB
DIP2C
PRKD1
TP53
NLRP10
GAB2
CDC42BPG
TGCT.perCancer.coldhotdiff.fisher
UCEC.perCancer.coldhotdiff.fisher
ASH1L
UCS.perCancer.coldhotdiff.fisher
FCGBP
UVM.perCancer.coldhotdiff.fisher
Cold.vs.hot.mut.frac
-log10(p.value)
pass
M
N
Y
direction
coldenrich
hotenrich
nochange

**Supplementary Figure S2. Cancer-specific relative enrichment of mutations in T cell-inflamed and non-inflamed tumors from all protein-coding genes on the genome.** 23 cancer types are shown here; 6 cancer types are shown in **Figure 2**. Somatic mutation data were not available for mesothelioma or primary melanoma tumors, hence not shown. Cold-enrich (yellow) represents genes with NSSMs enriched in non-T cell-inflamed tumors; hot-enrich (cyan) represents genes with NSSMs enriched in T cell-inflamed tumors. P-value shown as raw values without FDR-correction for multiple testing effect. Cancer ID and description can be found in **Supplementary Table S1**.

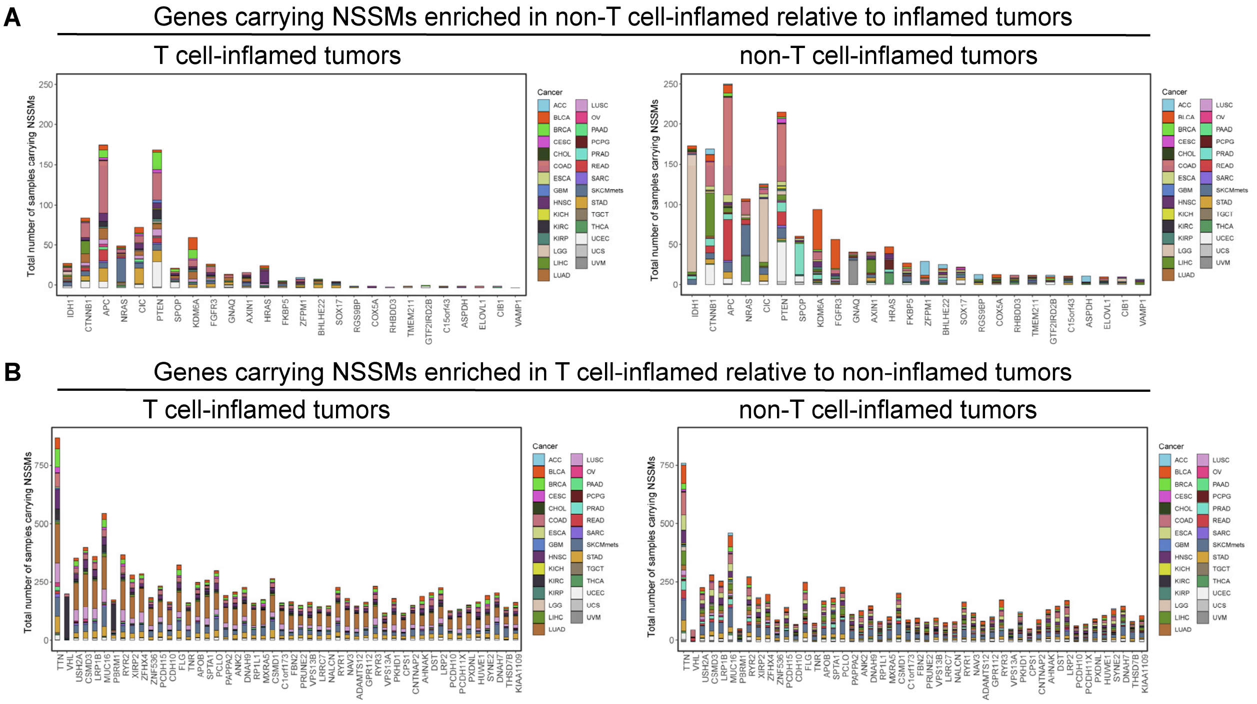


**Supplementary Figure S3. Pan-cancer distribution of mutations enriched in T cell-inflamed or non-inflamed tumors in all protein-coding genes on the genome, colored by cancer type.** (**A**) 26 genes carrying NSSMs more frequent in non-T cell-inflamed relative to inflamed tumors at FDR-corrected *P*<0.20. (**B**) Top 50 genes carrying NSSMs more frequent in T cell-inflamed relative to non-inflamed tumors at FDR-corrected *P*<0.05. Genes are listed on x-axis, and the number of tumors carrying NSSMs is shown on the y-axis. Each color represents individual cancer types. Cancer ID and description can be found in **Supplementary Table S1**.

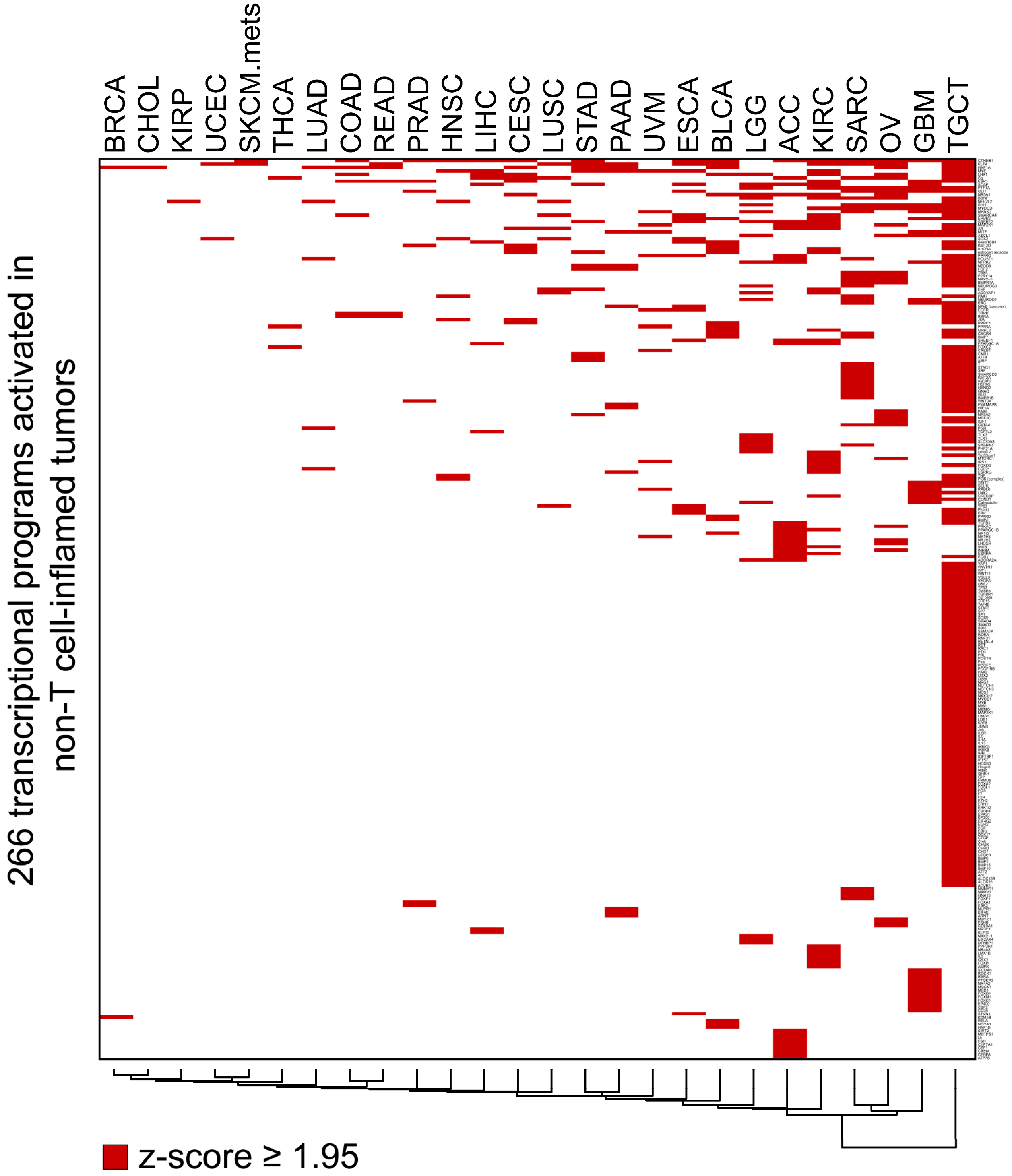


**Supplemnetary Figure 4. Pan-cancer pathway activation in non-T cell-inflamed relative to inflamed tumors from RNAseq gene expression, with all 266 pathways shown.** Each tumor type shows transcriptional programs activated in non-T cell-inflamed relative to the inflamed tumor group. Red color labels those at activation z-score ≥ 1.95 predicted by IPA causal network analysis (see **Methods**). List of the 266 pathways and their activation status in each cancer type is provided in **Supplementary Table S4**. Cancer ID and description can be found in **Supplementary Table S1**.

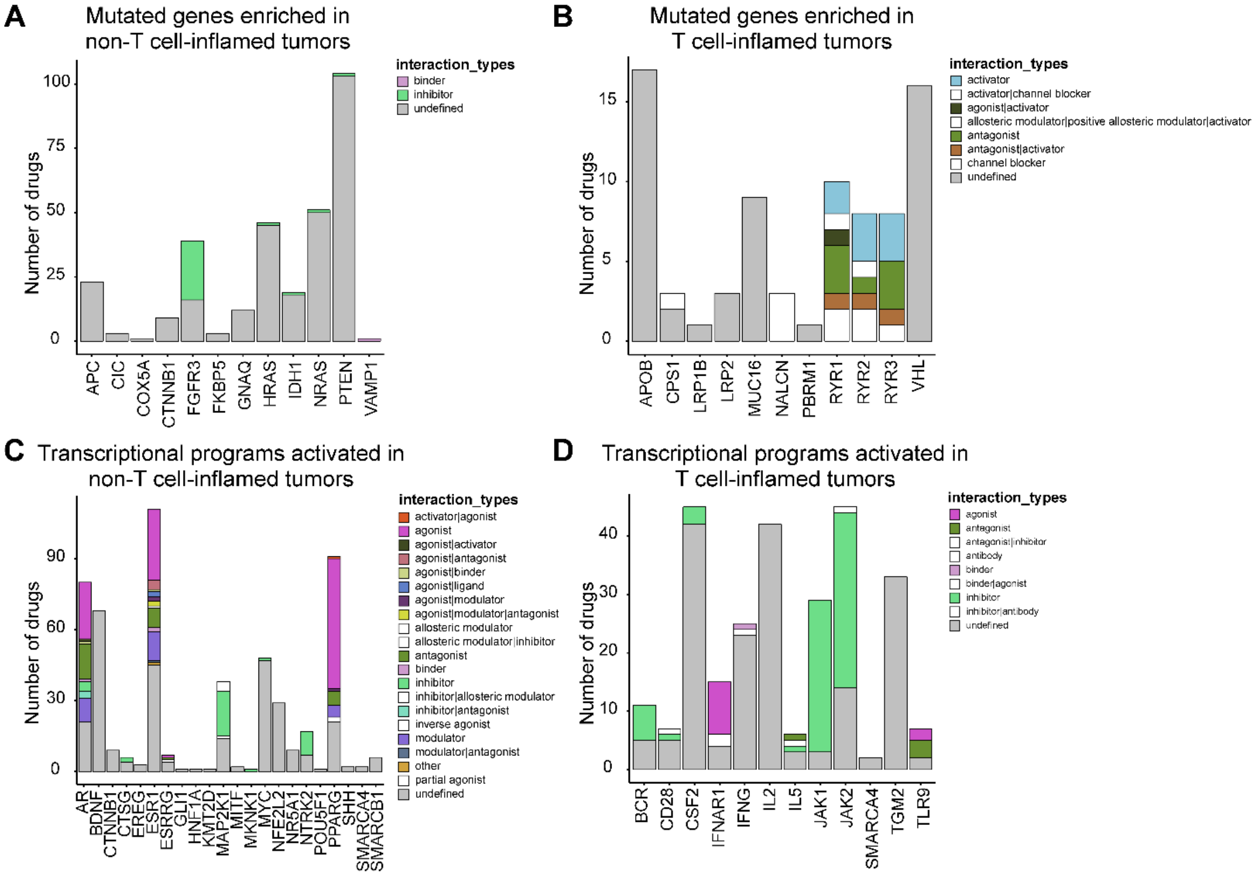


**Supplementary Figure S5. Summary of drug queries that target mutations enriched or transcriptional programs activated in non-T cell-inflamed and T cell-inflamed tumors.** (**A**) Drugs targeting the 26 cold-enrich mutated genes from **Figure 3A**. (**B**) Drugs targeting the 50 hot-enrich mutated genes from **Figure 3B**. (**C**) Drugs targeting the 31 transcriptional programs activated in non-T cell-inflamed tumors from **Figure 4A**. (**D**) Drugs targeting top 20 transcriptional programs activated in T cell-inflamed tumors. List of individual drugs for each gene or transcriptional program is provided in **Supplementary Table S6.**

### Supplementary Tables

**Supplementary Table S1.** List of cancer ID and description from TCGA included in this study.

**Supplementary Table S2.** List of genes investigated in individual cancer type analysis with their relative enrichment in the T cell-inflamed or inflamed phenotype. All genes are shown without filtering by p-values.

**Supplementary Table S3.** List of genes investigated in pan-cancer analysis with their relative enrichment in the T cell-inflamed or inflamed phenotype. All genes are shown without filtering by p-values.

**Supplementary Table S4.** List of pathways predicted to be activated in non-T cell-inflamed relative to inflamed tumors (n=266). Also shown in **Supplementary Figure S4**. 1 = activated; 0 = not-activated.

**Supplementary Table S5.** List of genes carrying NSSMs enriched in non-T cell-inflamed tumors (n=26, from **Figure 3A**) or transcriptional programs activated in non-T cell-inflamed tumors across at least 4 cancer types (n=31, from **Figure 4A**) at per patient level. For genes (columns denoted with “NSSMs”), 1 = mutated, 0 = not-mutated. For pathways (columns denoted with “pathways”), 1 = activated, 0 = not-activated.

**Supplementary Table S6.** List of drugs targeting relevant molecular mechanisms identified in our study from The Drug Gene Interaction Database (DGIdb).

### Supplementary Data Files

**Supplementary Data File S1. RPPA validation of activated transcriptional programs at protein level.** Pearson’s correlation was computed between the protein score of upstream regulator or target molecules (when available) and RNAseq expression of T cell-inflamed gene signature on the x-axis, with r and p-value shown above each individual plot. ConGene = concordant gene list, *alias* of the T cell-inflamed gene signature.
